# Genetic dissection of grafting traits reveals regulators of graft union formation in grapevine

**DOI:** 10.64898/2026.10.06.757043

**Authors:** Marilou Camboué, Marine Morel, Enric Zehraoui, Maria Lafargue, Cyril Hévin, Nicolas Hocquard, Jean-Pascal Tandonnet, Nathalie Ollat, Elisa Marguerit, Sarah Jane Cookson

**Affiliations:** EGFV, Univ. Bordeaux, Bordeaux Sciences Agro, INRAE, ISVV, F-33882, Villenave d’Ornon, Bordeaux France; Hennessy, rue de la Richonne – CS20020, 16101 Cognac cedex - France; INRAE, Univ. Bordeaux, UMR Biologie du Fruit et Pathologie, 1332, Villenave d’Ornon, France

**Keywords:** callus, grafting, healing, QTL, rootstock, scion, *Vitis* spp., wound

## Abstract

Understanding how grafts form has fascinated scientists for over a century, and numerous transcription regulators involved in wound healing, callus formation, and grafting have been identified in model plants. However, the genetic architecture underlying grafting traits remains poorly characterized. To address this gap, we aimed to identify quantitative trait loci (QTLs) associated with grafting success rates and grafting-related traits (including callus formation, mechanical strength of the graft union, and post-grafting growth) in a grapevine rootstock mapping population, *Vitis berlandieri × V. rupestris*. Although no QTLs were identified for grafting success rate, QTLs were identified for component traits revealing a genetic architecture that would have remained hidden when considering grafting success as a single composite phenotype. By integrating QTL and transcriptome data, we identified regulators potentially mediating grafting-related traits. We then showed that one transcription factor that regulates callus formation and grafting in Arabidopsis potentially has a similar role in grapevine by silencing with exogenous double-stranded RNAs and that exogenous application of methyl jasmonate increases callus development at the graft interface. Candidate transcriptional regulators located within QTLs and evidence from gene silencing and hormone treatments suggest the conservation of regulatory mechanisms underlying wound-induced callus formation across plant species.

**Highlight:** Grafting success is a complex phenotype that can be dissected into component traits. The combination of genetics, genomics and hormone treatments identified regulators controlling callus and potentially graft development.

## Introduction

Graft union formation is a complex and fascinating biological process. It begins with an initial wound response, followed by the exudation of pectin-like compounds that help bind the two partners. Subsequently, callus tissue proliferates, and, finally, the phloem and xylem connect, creating a mechanically resistant and functional union. Numerous studies have explored structural and anatomical changes during grafting, and many “omics” investigations have examined the accumulation of metabolites, proteins, and transcripts (reviewed in Loupit & Cookson (2020)). Not all scion/rootstock combinations graft together successfully and many show poor graft compatibility, i.e. the graft union that forms is weak with few vascular connections; these plants have poor growth and reduced survival. Identifying the regulators of graft compatibility is an important goal for the selection of new genotypes in grafted crop species.

Several genes regulating wound healing, callus formation, and graft union development have been identified, particularly in model species such as Arabidopsis. Transcription factors (TFs) are especially important. A well-known example is the *ETHYLENE RESPONSE FACTOR/APETALA2 (ERF/AP2) TF WOUND INDUCED DEDIFFERENTIATION1 (WIND1)*, which controls callus formation and petiole graft union development in Arabidopsis (Iwase *et al*., 2021). Other ERF TFs, particularly those of the X subgroup (ERF108–115), contribute to jasmonate signaling as well as wounding and regeneration responses (Heyman *et al*., 2018; Sui *et al*., 2025). For example, ERF115 forms a heterodimer with the GRAS TF PHYTOCHROME A SIGNAL TRANSDUCTION 1 (PAT1) to regulate tissue regeneration after wounding via WIND1 (Heyman *et al*., 2016). Recently, PAT1 was also shown to influence both callus formation and grafting success in Arabidopsis and Norway spruce (Feng *et al*., 2024b).

Compared with research on traits such as disease resistance or fruit quality, genetic studies of grafting success have advanced slowly. This is largely due to the difficulty of reliably phenotyping grafting success rates in large populations, which ideally requires many replicates. In perennial crops, producing sufficient woody stems or buds for large-scale experiments can take many years. As a result, researchers have explored proxies for grafting success, such as metabolites (Loupit & Cookson, 2020), anatomic descriptions on the graft interface (Irisarri *et al*., 2021; Thomas *et al*., 2023) or assessing the morphology of grafted plants (Tedesco *et al*., 2020). However, no metabolite marker with sufficient predictive power has been identified (Loupit & Cookson, 2020), and accurately characterizing anatomical differences at the graft interface remains challenging because it develops heterogeneously in three dimensions. Measuring the mechanical resistance of the graft union offers a potentially high-throughput, quantitative approach. This method has been applied to quantify adhesion strength in *in vitro* tobacco stem grafts (Kawakatsu *et al*., 2020), and the mechanical strength of grafts of a range of species (Lindsay *et al*., 1974; Melnyk *et al*., 2018; Roberts & Brown, 1961), but has not yet been used to compare different scion/rootstock combinations or in large scale genetics experiments.

Few genetic studies focused on grafting have been published. The genetics of graft incompatibility has been studied in Douglas fir, it was found that more closely related genotypes are more graft compatible and that variance in graft compatibility is probably controlled by many genes (Copes, 1973). Another study found that graft compatibility in Douglas fir is controls by additive genes with a heritability of 0.81 (Copes, 1974). Graft incompatibility in peach is thought to be due to two dominant alleles (Salesses & Al Kaï, 1985). More recently, the genetic control of graft incompatibility has been studied in an apricot scion population grafted onto plum rootstock (Irisarri *et al*., 2019; Pina *et al*., 2021). The first study reported genetic variation in anatomical traits, such as necrotic layer formation and tissue discontinuities at the graft interface (Irisarri *et al*., 2019). The second study identified six significant QTLs (only at the chromosome-level) for the extent of necrosis, wood discontinuities, and overall visual incompatibility symptoms, but was limited by a small sample size (92 individuals) and lack of heritability estimates (Pina *et al*., 2021). Our recent study investigated the genetic architecture of early growth traits in grapevine cuttings and grafts, including grafting success rates (Morel *et al*., 2026). Despite analyzing two scions grafted with a rootstock population over two years, we detected no significant QTLs for grafting success (Morel *et al*., 2026). Similarly, despite grafting 16,645 grapevine grafts, Munoz *et al*. (2026) only found limited scion genetic control of grafting success rates (Munoz *et al*., 2026). Although research on the genetic basis of grafting success remains scarce or of limited success, more progress has been made in characterizing the genetics of wound-induced callus formation (Htwe *et al*., 2024; Huang *et al*., 2021; Nguyen *et al*., 2020; Song *et al*., 2024; Zhang *et al*., 2020) and tissue regeneration (Lardon *et al*., 2020; Nagle *et al*., 2024), which have revealed multiple candidate genes.

The aim of this study was to identify potential regulators of grafting success in grapevine, an iconic perennial grafted fruit crop, using genetic and genomic approaches. Because detecting QTL of grafting success rate has proven challenging (Morel *et al*., 2026; Munoz *et al*., 2026), we developed methods to phenotype component traits underlying graft union formation. Firstly, we assembled a device to measure the mechanical resistance of the graft union, and secondly, we quantified callus production on isolated stem internode segments. We then characterized the genetic architecture of grafting success rates and the component traits mechanical resistance of the graft interface, callus formation (both at the graft interface and on isolated stem internode segments), and post-grafting growth. We used a grapevine rootstock mapping population (*Vitis berlandieri × V. rupestris*; Berl10585 × RupMartin1701, Morel *et al*. (2026)) grafted with the scion *V. vinifera* cv. Cabernet-Sauvignon (CS). This population was selected because it is the largest interspecific biparental population of grapevine with 449 individuals (Morel *et al*., 2026) and because commercial rootstocks with *V. berlandieri* and *V. rupestris* parents have variable grafting success rates (Carrere *et al*., 2022; Loupit *et al*., 2022). Because transcriptional regulation, cell wall modifications and hormone action are key to callus and graft formation in many species (Feng *et al*., 2024a; Loupit *et al*., 2023; Wang *et al*., 2024); we then focused on these gene categories. We identified the genes belonging to the categories transcriptional regulation, cell wall modifications and hormone action within the confidence intervals of the QTLs of grafting-related traits and up-regulated in response to grafting using data from our previously published resource (Loupit *et al*., 2025). Then, we tested the function of two candidate TFs regulating wound-induced callus formation in grapevine by applying exogenous double-stranded RNAs (dsRNAs). Finally, we determined the role of jasmonate signaling in graft union formation by exogenously applying methyl jasmonate to the graft interface.

## Materials and methods

### Origin of Plant Material

The Berl10585 × RupMartin1701 progeny used in this study was described previously (Morel *et al*., 2026). It was obtained by crossing the female *V. berlandieri* 10585 (BER) with the male *V. rupestris* cv. Martin (RUP), producing 449 F_ individuals in 2017. This progeny was planted in two experimental setups to produce plant material for experiments: one in a greenhouse (G) with individuals cultivated in 7 L pots (planted in 2017) and the second in a field (F) with individuals (planted in 2021 at INRAE Nouvelle-Aquitaine Bordeaux, France, 44°47′28″ N; 0°34′41″ W). Woody stem material from these setups is referred to as “Greenhouse” (G) and “Field” (F), representing two independent sources of plant material. Individuals were selected if they had sufficiently large stem diameter to be grafted (over approximately 6 mm) and had not shown rooting defects in previous experiments so that grafting success rates were not confounded with poor rooting (Morel *et al*., 2026). The scion genotype was *V. vinifera* cv. Cabernet-Sauvignon clone 412 (CS), grown in the same vineyard as the Field material. Dormant canes of the progeny were collected from the greenhouse in December 2023 and from the field in March 2024. Canes of CS were collected in January 2024 for grafting experiments and in January 2025 for the double-stranded RNA assay. All samples were stored at 4 °C until use.

### Grafting Technique

Prior to grafting, one-bud scions (CS) and de-budded rootstocks (Berl10585 × RupMartin1701 progeny) were prepared and rehydrated overnight. Wood offcuts were retained for the callogenesis capacity assays described below. Grafts were made with scions and rootstocks of approximately the same diameter using our standard protocol (Loupit *et al*., 2025). Briefly, grafts were assembled with a mechanical grafting tool fitted with an omega-shaped blade (Omega Star, Chauvin, France). The unions were coated in melted wax (Staehler Rebwaches Pro containing 0.0035 % dichlorobenzoic acid) and placed in plastic boxes. After a 2 d resting period at ambient temperature, the grafts were transferred to a stratification room at 28 °C for 20 d, with approximately 1 cm of a 40 mg L_¹ copper sulfate solution added to the bottom of each box to avoid any fungi contaminations. Following stratification, grafts made with G origin were planted in biodegradable pots on a heated bench (25 °C) inside a hoop tunnel greenhouse. Whereas grafts made with F origin were planted directly in a field nursery at INRAE Nouvelle-Aquitaine Bordeaux (44°47′28″ N; 0°34′41″ W).

### Graft development phenotyping

The mechanical resistance of the graft union was quantified 6 to 7 weeks post-grafting. To do this, we assembled a crank bench stand with two clamps to attach to the scion and rootstock. We continuously pulled apart the graft and measured the mechanical resistance of the graft interface to the breaking point with a SAUTER FS 2-500 force gauge (Sauter GmbH, Germany) (Fig. 1A). After breaking the graft interface, we removed callus tissue from the graft interface with a scalpel and weighed. Measurements were taken from 8–12 April 2024 for the origin G and 15–18 June 2024 for the origin F, with 5-10 and 5 grafts per combination assed for G and F respectively. Shoot dry biomass was measured for all grafts whereas only the number of roots was recorded for the origin G.

**Figure 1.**
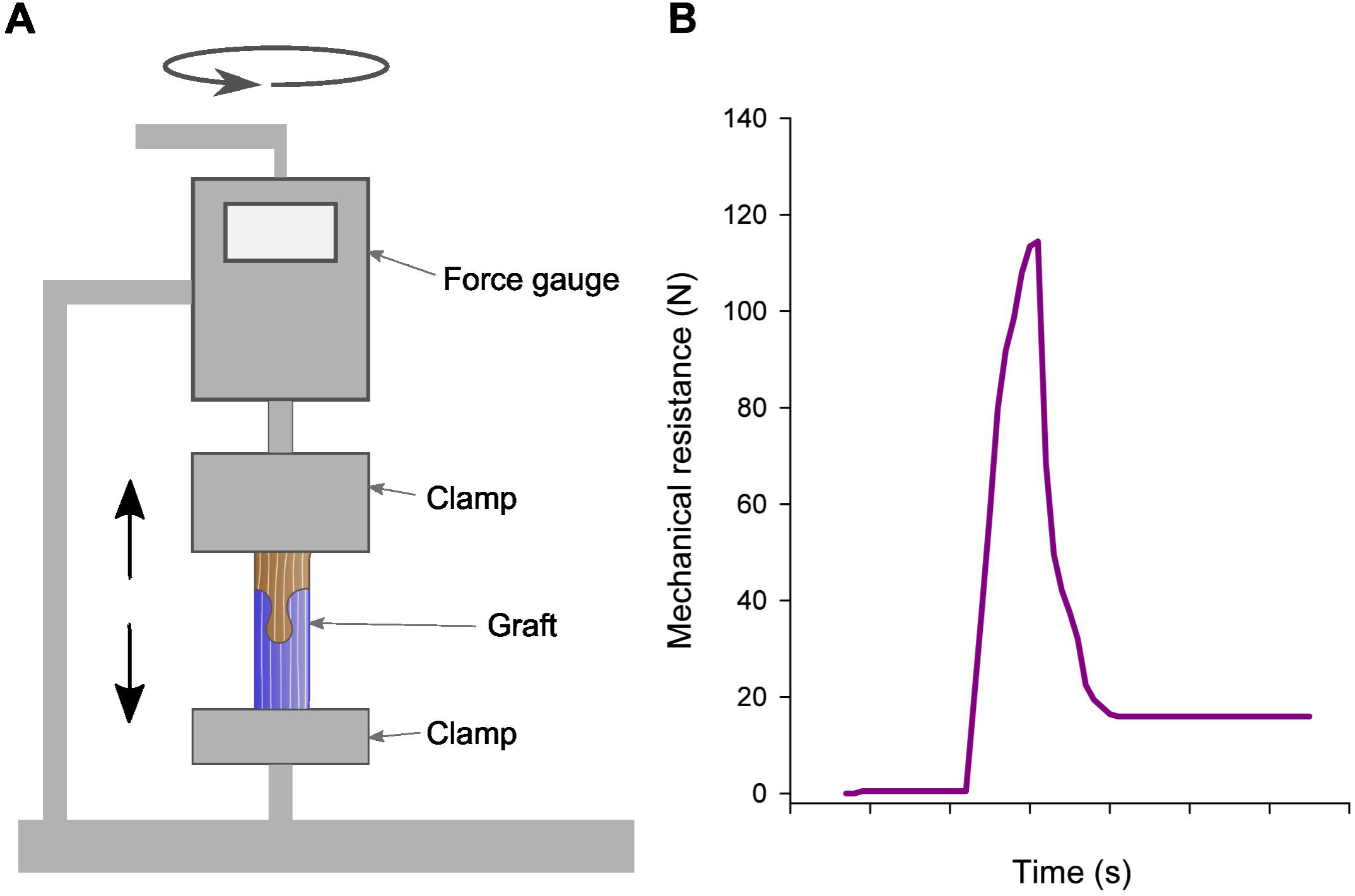
Measuring the mechanical resistance of graft union. A) Design of the device used to measure the mechanical resistance. Grafts were placed between two clamps, one connected to the base and the other connected to a force gauge. The grafts were pulled apart to breaking point by rotation of a handle. B) An example of the reading from the force gauge, the maximum value recorded before rupture was used to determine the maximum resistance of the graft union. ALT TEXT: A) Sketch of a device that measures the mechanical resistance of the graft interface, the scion and rootstock are attached to clamps with the scion clamp attached to a force gauge, the graft is then pulled apart and the force required to break the graft union is measured. B) An example of the readout from the force gauge showing changes with time in Newtons, which starts at 0 and increases to 120 Newtons before breaking.

The resistance of the graft interface to a manual “thumb test” was done on all remaining G and F grafts on 5 June 2024 and 4 December 2024 (corresponding to 120 and 225 d after grafting) respectively. The number of replicates varied from 3 to 10. Grafting success rates at “thumb test” for the progeny grafted with CS scions in 2021 and 2022 one have been previously published (Morel *et al*., 2026) and added to our analysis. Grafting success rates were calculated only for genotypes with at least three repetitions.

### Callogenesis capacity assays

The wood offcuts used for the callogenesis capacity assays originated from the same origins as those used for the graft development phenotyping, i.e. G and F. Two stem internode segments of different lengths (approximately 31 and 51 mm) were cut longitudinally with disinfected pruning shears. Each split internode segment was placed with the cut surface facing upwards on a moistened blotting paper (Fig. 2) corresponding to four split internode segments per genotype and plant material origin. The Petri dishes were incubated at 28 °C in darkness, reproducing stratification conditions for 20 d. Various traits were studied on each fragment, such as, presence/absence of callus, time to the first visible callus development and callus biomass after 20 d of development.

**Figure 2.**
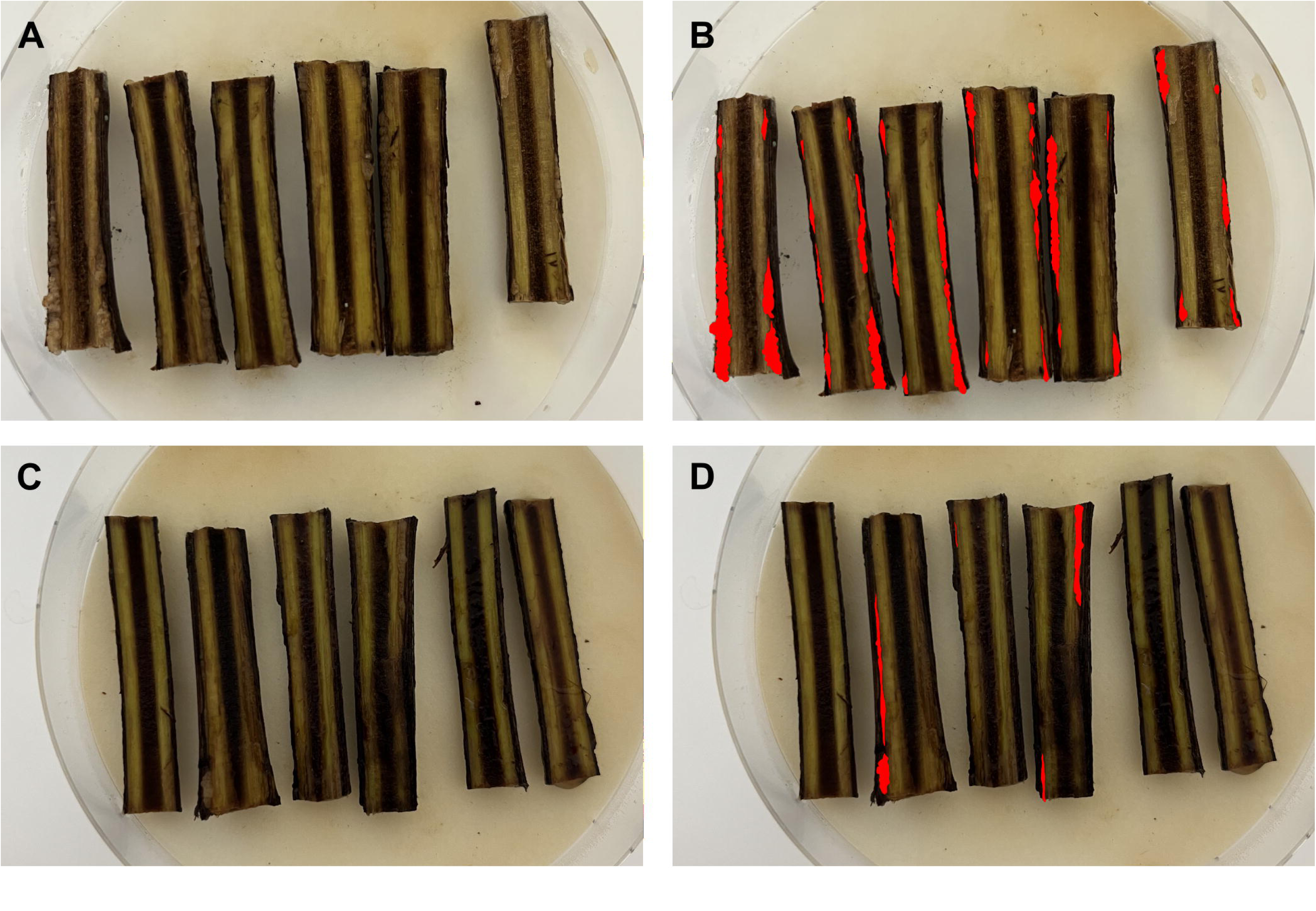
Callus surface area measurement on split internode segments treated with dsRNAs. dsRNAs designed to target either *GREEN FLUORESCENT PROTEIN* (A, B) or *PAT3/Vitvi10g00271* (C, D). Images taken 14 d after dsRNA treatment, (A, C) raw images, (B, D) images with segmentation of callus tissue area (shown in red) that was measured in imageJ. ALT TEXT: Photographs of petri dishes containing six split internode segments with the wounded side face up, segments are on humid filter paper with some callus development is visible along the damaged cambial tissues. Images taken 14 d after dsRNA treatment, dsRNAs designed to target either *GREEN FLUORESCENT PROTEIN* (A, B) or *PAT3/Vitvi10g00271* (C, D). (A, C) raw images, (B, D) images with segmentation of callus tissue area (shown in red) that was measured in imageJ.

### Number of genotypes studied in each experiment

For graft development phenotyping, 171 and 151 graft combinations were made from the progeny originating from the G and F respectively with 148 genotypes in common (Table S1). For the callogenesis capacity assays, 163 and 154 genotypes of the progeny from the G and F were phenotyped with 143 in common (Table S1). Overall, 168 genotypes were phenotyped for both graft development and callogenesis capacity (Table S1).

### Quantitative trait loci mapping and statistical analysis

All statistical analysis was done using R v4.2.0 statistics environment (R Core Team, 2014) using Rstudio (RStudio Team, 2020). Our data were analyzed separately according to the plant material origin, treating them as two independent datasets. For each trait, descriptive statistics (mean, minimum, maximum, standard deviation) and broad sense heritability (as genetic variance by sum of genetic and residual variance from the random model) were calculated. The difference between the two plant material origins was tested by variance analysis with aov(). Correlation between traits were calculated and plotted using cor() from ‘stats’ package and visualization with ggcorrplot() (Kassambara, 2022) from ‘ggplot2’package (Wickham, 2016).The QTL analyses for each trait were performed using BLUPs extracted from the linear random-effects model with rootstock genotypes as a random effect with the lmer() function from ‘lme4’ package (Bates *et al*., 2015) on the two parental genetic maps from Morel *et al*. (2026). A single QTL genome scan was carried out by Interval Mapping with the scanone() function from ‘R/qtl’ package (Broman *et al*., 2003). For each trait, the Logarithm of the Odds/LOD threshold (significance level α = 5%) was determined using 1000 permutations, both at the whole-genome and chromosome levels. Due to the high number of chromosome-level QTLs detected, an additional filtering criterion was applied to retain only high-confidence QTLs. Only QTLs with a LOD score above the significance threshold and a LOD score greater than 2 were retained. The peak LOD score was used to define the position of each QTL. For each QTL, the proportion of variance explained was estimated using the fitqtl() function. The approximate Bayesian credible interval (95%) was calculated to define the confidence interval of the QTL position. Gene positions within the interval boundaries were also recorded. In addition, the effect of each allele was evaluated by comparing the mean phenotypic values of individuals that were homozygous or heterozygous at the corresponding locus. When multiple peaks were detected within the same linkage group, their presence was further assessed using Composite Interval Mapping and multiple-QTL models were tested as done Morel *et al*. (2026).

### Wounding and grafting responses of genes within QTLs

Genes present within the confidence interval of most interesting QTLs were identified and their expression profile in response to wounding and grafting studied in our publically available dataset (Loupit *et al*., 2025). This dataset used describes the transcriptome of intact, wounded and homografted scions (with buds) and rootstocks (without buds) 14 d after wounding/grafting in *V. vinifera* cv. Pinot noir. The transcriptional regulators, cell wall and hormone related genes were identified based on their assignment to the MapMan BINS “RNA biosynthesis transcriptional regulation”, “Cell wall organization” and “Phytohormone action” respectively (Schwacke *et al*., 2019). Genes were selected with a Log2 fold change >1 and false discovery rate corrected p value <0.05 and expression in every condition studied. Log2(TMM) (Trimmed Mean of M component) normalized values were used to make HeatMaps using the pHeatmap (Kolde, 2025) package.

### Exogenous dsRNA application

Two candidate TFs involved in callus formation were selected from our data and the literature (*PAT3/Vitvi10g00271* and *ERF115/114/Vitvi17g00025*) along with one control gene not present within the grapevine genome (*GREEN FLUORESCENT PROTEIN, GFP*). Gene specific amplicons ∼250 bp were amplified from either CS wood cDNA or gDNA. For cDNA synthesis, RNA extraction and cDNA synthesis was performed as described in Cookson *et al*. (2013) and gDNA was extracted using the protocol described by Berger *et al*. (2022). The plasmid pGFPGUSPlus was used for the GFP sequence (addgene, plasmid pGFPGUSPlus #64401). A T7 promoter sequence was added to the 5′ end of each primer to enable in vitro transcription. Amplicons were purified using the Wizard® SV Gel and PCR Clean-Up System (Promega). *In vitro* transcription of dsRNA was performed with the HiScribe™ T7 Quick Kit (NEB) from 1 µg of purified template, with the NTP concentration increased to 5 µL to optimize yield. Reactions were incubated for 16 h at 37 °C. Synthesized dsRNA was purified by ethanol precipitation (0.1 vol. sodium acetate 3 M pH 5.5 + 2.5 vol. 100 % ethanol), washed with 70 % ethanol, air-dried, and re-suspended in 50 µL RNase-free water. Primers and sequences are provided in Table S2.

Stem internode segments (4 cm long) of one genotype (CS) were split longitudinally and 11 split segments were assigned randomly to each treatment. Treatments were dsRNAs targeting the control gene *GFP* and the grapevine genes *ERF115/114* and *PAT3*, as well as a combined treatment with *ERF115/114 + PAT3*. Split internode segments were rehydrated by being placed cut-side down in dsRNA solution (20 ng/µL) for 4 h at room temperature. Then these segments were placed with the inner surface facing up on moistened blotting paper in Petri dishes (5 or 6 segments per petri dish). Subsequently, 200 µL of dsRNA at 200 ng/µL was applied by pipette to each split internode segment at 1, 6, 11, 14, 16, 18, 21, 24 and 26 d after start of the experiment. Once callus tissue became visible, photographs were taken at regular intervals. Callus surface area was measured manually on the photographs using Fiji (ImageJ 1.54f, Java 1.8.0) (Fig. 2), from the first apparition of callus to the production of adventitious roots.

### Exogenous methyl jasmonate application

To test the effect of exogenous methyl jasmonate application on graft union formation, auto-grafts of CS were made (i.e. stems were grafted with themselves). The grafts were prepared with an omega grafting machine, disassembled and then the cut surfaces of the scion and rootstock were dipped into solutions containing either water or 100, 300 or 600 µM of methyl jasmonate for 5 min. The grafts were then reassembled, dipped in wax, stratified and planted in a hoop greenhouse as described above. Six weeks after grafting, the mechanical resistance of the graft interface and callus biomass was measured in 30 grafts of each treatment and significance was tested with an ANOVA on Ranks (SigmaPlot 16).

## Results

Grafts of CS scions onto the Berl10585×RupMartin1701 rootstock progeny were made with rootstocks originating from two different growing conditions, the field (F) and the greenhouse (G) allowing us to do two independent experiments in 2024. These grafts were phenotyped for various traits related to grafting success, such as, callus formation at the graft interface, the mechanical resistance of the graft union (using the equipment shown in Fig. 1) and post-grafting growth. We also assessed intrinsic callogenesis capacity on isolated stem fragments (as shown on Fig. 2).

### Grafting success rate varies from year to year

Grafting success rate after one season in a field nursery was also measured in this study (in 2024) and this data was analyzed in parallel with our previously published results from 2021 and 2022 (Morel *et al*., 2026). Grafting success rates varied considerably between years, ranging from 36% to 60%, with 154, 203, and 271 individuals assessed in 2024, 2021, and 2022, respectively (Table S1). 78 rootstock individuals were common to all years, these individuals gave a good representation of all the genotypes studied, but there was little stability in grafting success rates across the three years of study (Fig. S1).

### QTL detection of grafting related traits

The Berl10585×RupMartin1701 progeny showed considerable phenotypic variation in grafting success rates and measurements of grafting-related traits with coefficients of variation from 40 to over 200 % (Table 1). The distributions generally did follow a normal distribution according to the Shapiro–Wilk test for normality. The origin of the plant material significantly affected the mass of callus produced at the graft interface and in the callogenesis-capacity assays on split internode segments, but not the mechanical resistance of the graft interface and shoot biomass (Table 1). The number of roots was only measured in the greenhouse experiment because of the difficulty of recovering root systems in field conditions (Table 1). The broad sense heritability ranged from 0.29 for shoot biomass to 0.71 for callus biomass (Table 1). As there was a significant difference between the development of grafts and callus on internode segments made with plant material of different origin, i.e. F and G origins. As a consequence, the data from the F and G experiments were analyzed separately for correlations between traits and subsequent QTL detection.

**Table 1:** Summary statistics of the key phenotypic data and comparison between plant material originating from the field and greenhouse.

| **Trait** | **Plant material origin** | **Min** | **Max** | **Mean** | **Standard deviation** | **Coefficient of variation** | **p-value (T-test)** | **Heritability** |
| --- | --- | --- | --- | --- | --- | --- | --- | --- |
| **Graft union phenotypes** |  |  |  |  |  |  |  |  |
| **Callus fresh biomass (g)** | Field | 0 | 1.58 | 0.20 | 0.24 | 0.12 | 1.92e-05 | 0.34 |
|  | Greenhouse | 0 | 1.44 | 0.25 | 0.20 | 0.08 |  | 0.50 |
| **Mechanical resistance of the graft union (N)** | Field | 2.5 | 337 | 69.6 | 43.1 | 62.0 | 0.2103 | 0.38 |
|  | Greenhouse | 5 | 220 | 71.8 | 29.3 | 40.9 |  | 0.35 |
| **Shoot biomass (g)** | Field | 0 | 12.3 | 0.96 | 1.71 | 178.9 | 0.7423 | 0.31 |
|  | Greenhouse | 0 | 10 | 0.92 | 0.65 | 70.3 |  | 0.29 |
| **Number of roots** | Greenhouse | 0 | 28 | 5.9 | 4.1 | 68.9 |  | 0.44 |
| **Callogenesis capacity phenotypes** |  |  |  |  |  |  |  |  |
| **Callus fresh biomass (g)** | Greenhouse | 0 | 0.54 | 0.03 | 0.06 | 205.56 | 2.424e-14 | 0.71 |
|  | Field | 0 | 0.99 | 0.08 | 0.14 | 179.91 |  | 0.51 |

*Comparison of traits between grafts made with plant material from the field and greenhouse tested with a T-test.*

We found that grafting success rate (i.e. percentage of grafts that had graft unions resistant to breaking under the “thumb test”) was significantly correlated with shoot biomass and the mechanical resistance of the graft interface in both the F and G experiments (Fig. 3). In the F experiment, grafting success rate was also correlated with the mass of callus produced at the graft union (Fig. 3B). However, none of the traits studied in the callogenesis capacity assays on isolated internode segments were correlated with the grafting success, but a significant correlation was found between callus fresh weight on isolated stem internode segments and callus dry weight at the graft interface (Fig. 3). There was a negative correlation between the time to produce the first visible callus cells and callus biomass (Fig. 3).

**Figure 3.**
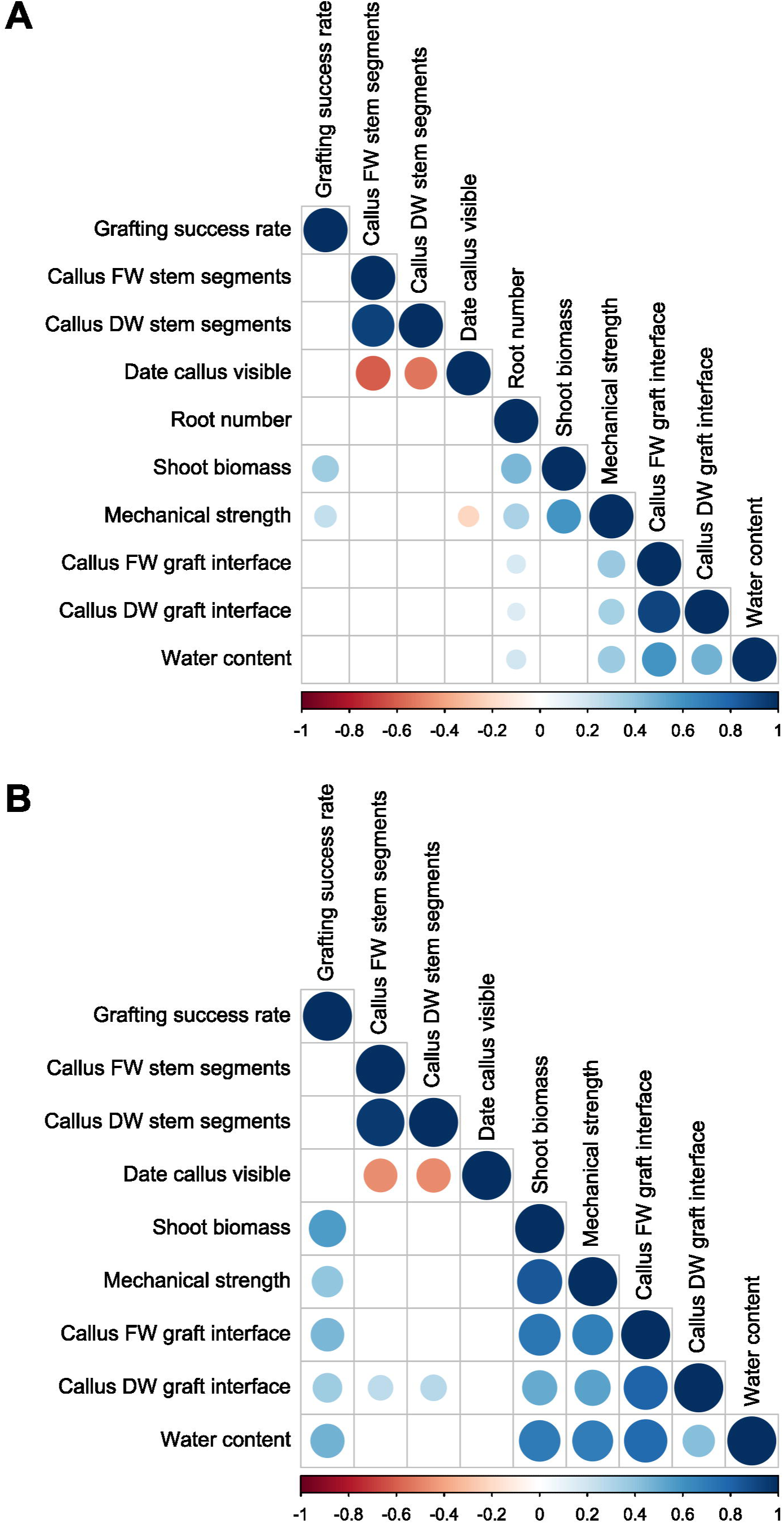
Significant Pearson’s correlation coefficients between grafting-related traits. Plant material from A.) the greenhouse and B.) the field. ALT TEXT: Correlation matrixes of significant Pearson’s correlation coefficients for plant material from A.) the greenhouse and B.) the field. Positive correlations found between most grafting-related traits.

We were unable to detect significant QTL for both grafting success rates (despite quantifying grafting success in three separate experiments on 203, 271 and 154 genotypes) and callus formation on isolated stem internode segments. However, the QTL analysis identified many grafting-related QTLs of component traits explaining a small percentage of phenotypic variance (generally below 10 %) (Table 2). Six QTLs were significant at the whole genome level and 13 QTL were significant at the chromosome level (Table 2). Some QTLs are of particular interest (i.e. most likely to be related to graft union formation such as mechanical resistance of the graft union and callus biomass) and were found on LG1 LG7, LG9, LG10, LG13, and LG19.

**Table 2.** Quantitative trait loci identified for grafting-related traits. The plant material used had two different origins (greenhouse (G) and field (F)) and two parental maps were used, *Vitis berlandieri* (BER) and *V. rupestris* (RUP).

| Plant material origin | Map | LG | Position (cM) | logarithm of the odds (LOD) score | LOD threshold 5 % on LG | LOD threshold 5 % whole genome | Marker closest to peak | SNP position in cM | Percentage of explained variance | Left flanking marker | 95 % confidence intervals (cM) | Right flanking marker | aa/ab |
| --- | --- | --- | --- | --- | --- | --- | --- | --- | --- | --- | --- | --- | --- |
| **Callus biomass graft interface** |  |  |  |  |  |  |  |  |  |  |  |  |  |
| F | BER | 7 | 57.5 | 2.4 | 1.8 |  | chr07_22436689 | 57.5 | 7.5 | chr07_6739260 | 28.6-73 | chr07_27327459 | 171./263.7 |
|  | BER | 10 | 17.5 | 2.4 | 1.8 |  | chr10_3499321 | 17.5 | 7.7 | chr10_1353125 | 7-39 | chr10_7412036 | 159.5/254.7 |
|  | BER | 1 | 54.7 | 2.7 | 1.7 |  | chr01_22976412 | 54.7 | 8.6 | chr01_360889 | 0-63.2 | chr01_23937167 | 168.3/265.7 |
|  | RUP | 13 | 64.2 | 2.1 | 1.8 |  | chr13_22480071 | 64.2 | 6.6 | chr13_2816105 | 16-72.0 | chr13_26254495 | 167.47/266.32 |
|  | RUP | 9 | 8 | 3.0 |  | 3.0 | chr09_1097398 | 7.6 | 9.2 | chr09_243351 | 0.91-23 | chr09_4761527 | 154.6/263.1 |
| G | RUP | 13 | 8.4 | 2.3 | 1.8 |  | chr13_1587618 | 8.4 | 5.8 | chr13_623448 | 0-83.6 | chr13_29028302 | 189.2/268.5 |
|  | RUP | 19 | 4.9 | 2.3 | 1.7 |  | chr19_1733249 | 4.9 | 6.7 | chr19_49116 | 0-41.1 | chr19_7676767 | 188.9/266.6 |
|  | RUP | 7 | 37.6 | 2.6 | 1.9 |  | chr07_7882093 | 37.6 | 8.5 | chr07_2607868 | 17-71 | chr07_19406858 | 177.8/274.9 |
| **Mechanical resistance of the graft union** |  |  |  |  |  |  |  |  |  |  |  |  |  |
| F | BER | 7 | 57.5 | 3.1 |  | 2.9 | chr07_22436683 | 57.5 | 9.9 | chr07_21492933 | 54.0-62.2 | chr07_23769013 | 60.9/80.4 |
| G | BER | 11 | 10.7 | 2.1 | 1.7 |  | chr11_1870356 | 10.7 | 6.0 | chr11_417738 | 0-41.7 | chr11_16111163 | 65.9/76.0 |
| **Shoot biomass** |  |  |  |  |  |  |  |  |  |  |  |  |  |
| F | RUP | 18 | 81.0 | 2.2 | 1.8 |  | chr18_21328594 | 80.6 | 6.7 | chr18_9621616 | 53-114.092 | chr18_28114543 | 0.699/1.13 |
|  | BER | 7 | 57.5 | 3.3 |  | 2.8 | chr07_22436689 | 57.5 | 10.1 | chr07_16622804 | 38 / 70.26 | chr07_26417984 | 0.619/ 1.35 |
| G | RUP | 3 | 48.8 | 3.1 |  | 3.0 | chr03_7330370 | 48.8 | 7.0 | chr03_6642923 | 46.00 - 64.66 | chr03_20430234 | 0.789/ 1.04 |
|  | BER | 11 | 10.7 | 3.6 |  | 3.0 | chr11_1870356 | 10.7 | 4.1 | chr11_417738 | 0-18.8 | chr11_3306467 | 0.874/ 0.926 |
|  | RUP | 17 | 60.3 | 2.7 | 1.8 |  | chr17_14963896 | 60.3 | 7.0 | chr17_2912855 | 10-62.27 | chr17_16553101 | 0.794/1.00 |
|  | BER | 7 | 61.4 | 4.0 |  | 3.0 | chr07_23769039 | 61.4 | 3.2 | chr07_19625792 | 49-66 | chr07_25378031 | 0.780/1.05 |
| **Number of roots** |  |  |  |  |  |  |  |  |  |  |  |  |  |
| G | BER | 11 | 9.5 | 2.1 | 1.6 |  | chr11_1595782 | 9.5 | 4.0 | chr11_417738 | 0-18.8 | chr11_3306467 | 5.13 / 6.68 |
|  | RUP | 17 | 56.2 | 2.3 | 1.7 |  | chr17_11216365 | 56.2 | 6.0 | chr17_2912855 | 10-62.27 | chr17_16553101 | 5.22/ 6.73 |
|  | BER | 1 | 63.0 | 2.5 | 1.7 |  | chr01_23945947 | 63.0 | 7.1 | chr01_360889 | 0-63.24 | chr01_23937167 | 5.29/ 6.74 |

### Identification of genes located within QTLs are up-regulated in response to grafting

The genes located within QTL intervals of interest were identified, and their expression patterns were examined using a transcriptome dataset of intact, wounded, and grafted scions and rootstocks of grapevine (Loupit *et al*. (2025); Tables S4, S5). Particular attention was given to the functional categories known to be involved in graft union formation, namely transcriptional regulation, cell wall organization and hormone action with a focus on those genes up-regulated in both scion and rootstock in response to grafting (Tables S4, S5; Fig. 4, Fig. S2).

**Figure 4.**
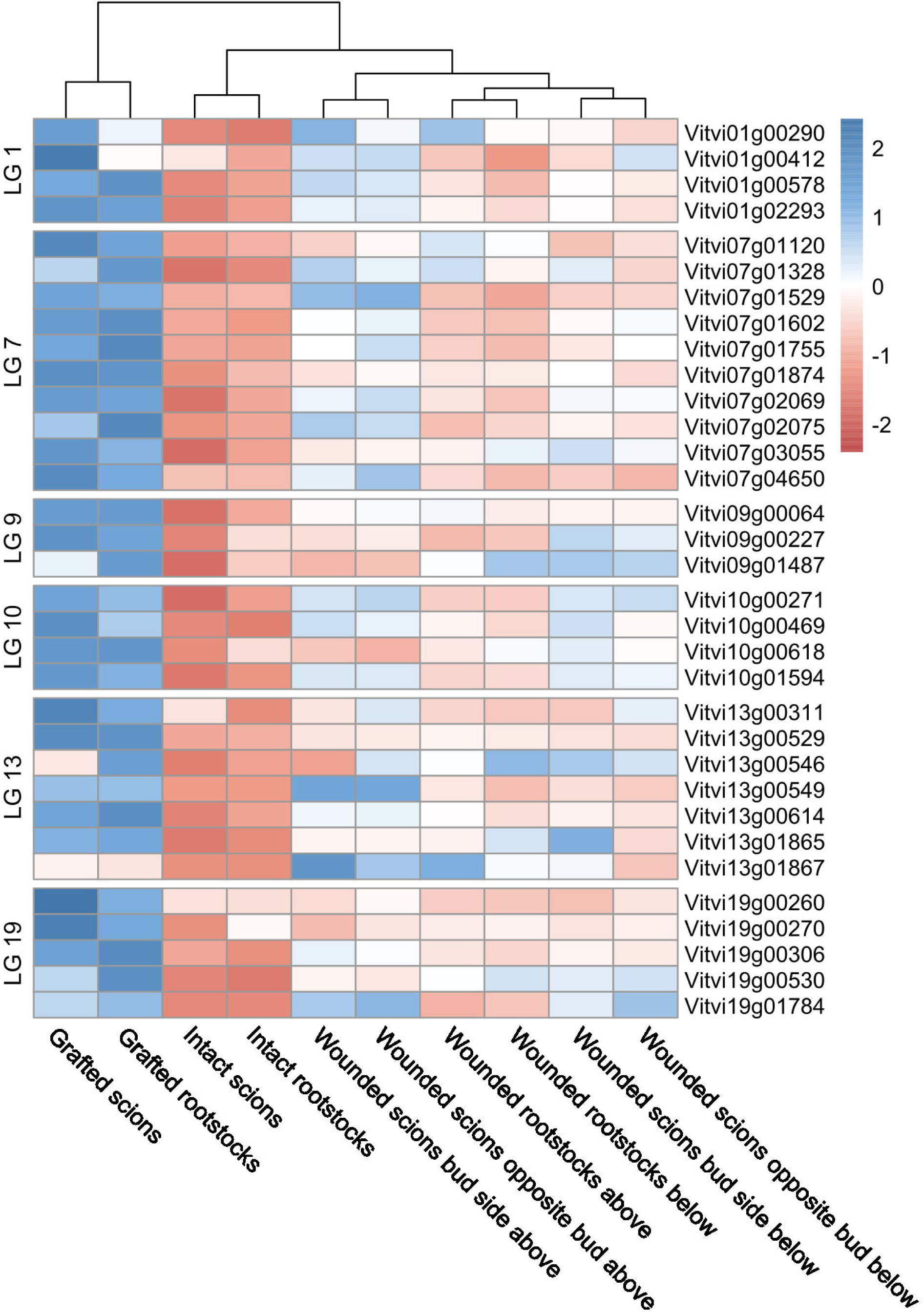
Heatmap of the expression levels of key transcriptional regulators in response to grafting. Expression of transcriptional regulators up-regulated in the scion and rootstock in response to grafting and within the confidence intervals of QTLs; expression shown in intact, wounded and grafted scion and rootstocks of grapevine 14 d after grafting (data from Loupit et al. (2025)). ALT TEXT: Heatmap showing the up-regulation of the expression of 33 transcriptional regulators located within QTLs on linkage groups/LGs 1, 7, 9, 10, 13 and 19. Lowest expression is found in intact scions and rootstocks, and highest expression is found in grafted scions and rootstocks, wounded scions and rootstocks have intermediate expression levels.

On LG1, two overlapping QTLs significant at the chromosome level were detected: one for callus biomass and another for root number (Table 2). Although their peak positions differed, the confidence intervals overlapped, spanning 1,586 genes. Of these, 81 were up-regulated in both scions and rootstocks (Table S4). Four transcriptional regulators were among them: the transcriptional repressor *JAZ10 (Vitvi01g02293)*, *LBD42 (Vitvi01g00290)*, a homeobox-leucine zipper protein (*Vitvi01g00412*), and a MYB (*Vitvi01g00578*) (Table S5).

On LG7, QTLs were associated with callus biomass, mechanical resistance at the graft interface, and shoot biomass. The QTLs for callus biomass were significant at the chromosome level, whereas the others were significant at the whole-genome level (Table 2). The confidence interval contained 1,324 genes, of which 61 were up-regulated in both scions and rootstocks after grafting (Table S4). Among these, 10 were transcriptional regulators (Fig. 4), including three ERF109/110 members, two NAI1 orthologues, two MYBs (a MYB93, *Vitvi07g02075* and *MYB148/Vitvi07g02075*) and one transcriptional adaptor, the NINJA family protein *Vitvi07g01120* (Table S5).

On LG9, a genome-wide significant QTL for callus biomass was detected (Table 2). This interval contained 439 genes, 14 of which were up-regulated in both graft partners (Table S4). Three of these were transcriptional regulators: *bHLH87* (*Vitvi09g00227*), *GRAS/NSP1* (*Vitvi09g01487*), and the transcriptional repressor *JAZ6* (*Vitvi09g00064*) (Table S5).

On LG10, a QTL significant at the chromosome level was associated with callus biomass at the graft interface (Table 2). This interval contained 672 genes, with 45 up-regulated in both scion and rootstock in response to grafting (Table S4). Only four were TFs: *HEAT STRESS FACTOR B-2A* (*Vitvi10g00469*), an ERF (*Vitvi10g01594*), *PAT3/Vitvi10g00271*, and *WRKY65* (*Vitvi10g00618*) (Table S5).

On LG13, two overlapping QTLs significant at the chromosome level were identified for callus biomass from both greenhouse and field repetition (Table 2). The smaller interval contained 1,107 genes, of which 45 were up-regulated in response to grafting (Table S4). These included seven TFs: three LBD11 genes, one ERF (*TARGET OF EARLY ACTIVATION TAGGED 2*/*TOE2/Vitvi13g00529*), one *GATA* (*Vitvi13g00614*), and two *GRAS SCL PROTEINS* (*SCL14/Vitvi13g00311* and *SCL9/Vitvi13g01865*) (Table S5, Fig. 4).

On LG19, a QTL significant at the chromosome level was associated with callus biomass (Table 2). Its confidence interval included 638 genes, with 31 up-regulated in both scions and rootstocks (Table S4). Five TFs were identified: two ERFs, *MYB102* (*Vitvi19g00306*), *NAC72/JA2L* (*Vitvi19g00270*), and a *WRKY* (Table S5).

In addition to transcriptional regulators, genes involved in cell wall organization and hormone action were also studied (Fig. S2). The eight genes from the category cell wall organization covered all aspects of cell wall biology including cellulose synthesis, cutin/suberin formation, hemicellulose formation, pectin synthesis and Casparian strip formation (Fig. S2A). Thirteen genes associated with hormone action were identified (Fig. S2B). These included one gene involved in abscisic acid transport (*Vitvi01g00921*); three genes involved in auxin conjugation and degradation (*Vitvi01g02092*, *Vitvi19g00369* and *Vitvi19g00574*); one gene involved in cytokinin biosynthesis (*Vitvi01g00298*); and two cytokinin dehydrogenases located on chromosome 7 (*Vitvi07g04213* and *Vitvi07g04222*), which were more strongly upregulated in the rootstock than in the scion. Two genes involved in jasmonic acid biosynthesis were also identified, encoding *LIPOXYGENASE 3* (*LOX3*/*Vitvi09g00085*) and an allene oxide cyclase (*Vitvi01g00246*), together with JAZ6 (*Vitvi09g00064*), a gene involved in jasmonic acid signaling that encodes the transcriptional repressor described above. In addition, three signal peptide genes belonging to the non-cysteine-rich peptide category were identified (*Vitvi10g01798*, *Vitvi01g01265* and *Vitvi01g01924*).

### Exogenous application of dsRNAs targeting TFs potentially regulating callus formation

Potential regulators of callus formation were selected from the literature, and our QTL and transcriptomic data. *PAT3* was selected because it was located within a QTL, was up-regulated in response to grafting and orthologues of *PAT3* have been shown to regulate graft union formation in multiple species (Feng *et al*., 2024b). As PAT1 interacts with ERF115 in Arabidopsis (Heyman *et al*., 2016), the combined silencing of *PAT3* and the grapevine orthologue of *ERF115* was also tested. There was a significant reduction in the surface area of callus formed in response to wounding after the addition of dsRNAs targeting *PAT3* and *PAT3 + ERF* in comparison to the control (*GFP*) from 14 d after wounding (Fig. 5). The application of dsRNAs designed to target *PAT3* also delayed the appearance of callus on isolated stem internode segments (Fig. 5B). We found that the surface area of callus was significantly correlated to callus fresh weight (Pearson correlation coefficient of 0.967 and P value 1.437E-57, Fig. S3).

**Figure 5.**
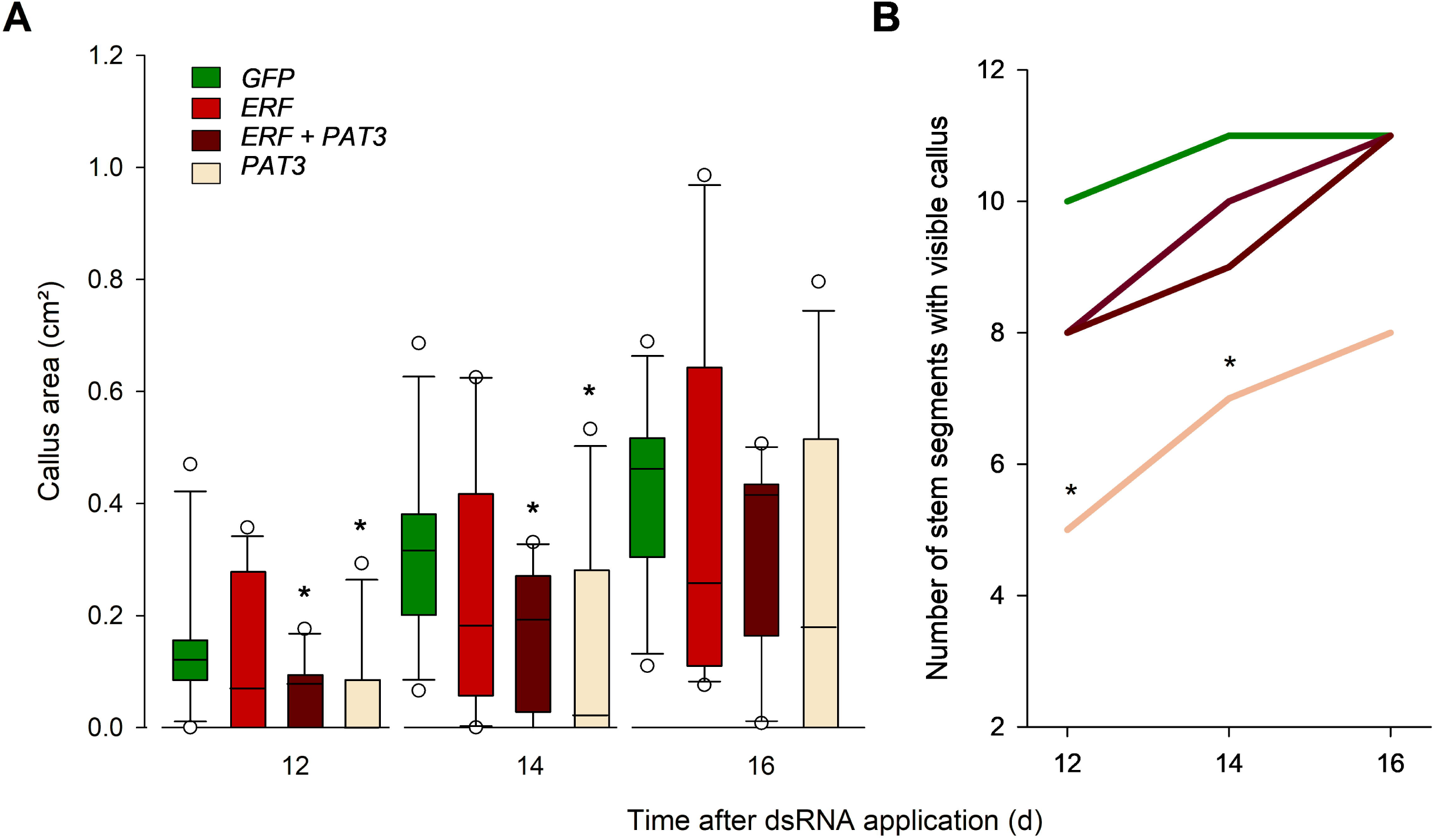
The effect of dsRNA application on A) callus area and B) visible callus appearance. dsRNAs were designed to target the transcription factors *PAT3/Vitvi10g00271*, *ERF/Vitvi17g00025* and PAT3 + ERF, dsRNAs targeting *GREEN FLUORESCENT PROTEIN/GFP* were used as a control (n = 11). Significant differences between *GFP* and transcription factor treatments indicated by stars, A) Mann-Whitney U test, B) z-test. ALT TEXT: A) Box plots of callus surface area measured on photographs (as shown in Figure 2) of split internode segments treated with different dsRNAs. B) Line graph of the number of internode segments with visible callus at 12, 14 and 16 d after dsRNA treatment.

### Exogenous application of methyl jasmonate stimulates callus formation

Because three transcriptional regulators and two jasmonate biosynthesis genes were found within the intervals of the QTLs and up-regulated in response to grafting, we tested the role of jasmonate in graft union formation. Exogenous application of methyl jasmonate at the graft interface immediately after grafting trigger an increase in callus biomass 6 weeks after grafting (Fig. 6A, tested with a Kruskal-Wallis One Way Analysis of Variance on Ranks, P = 0.044), but did not affect the mechanical resistance of the graft interface (Fig. 6B).

**Figure 6.**
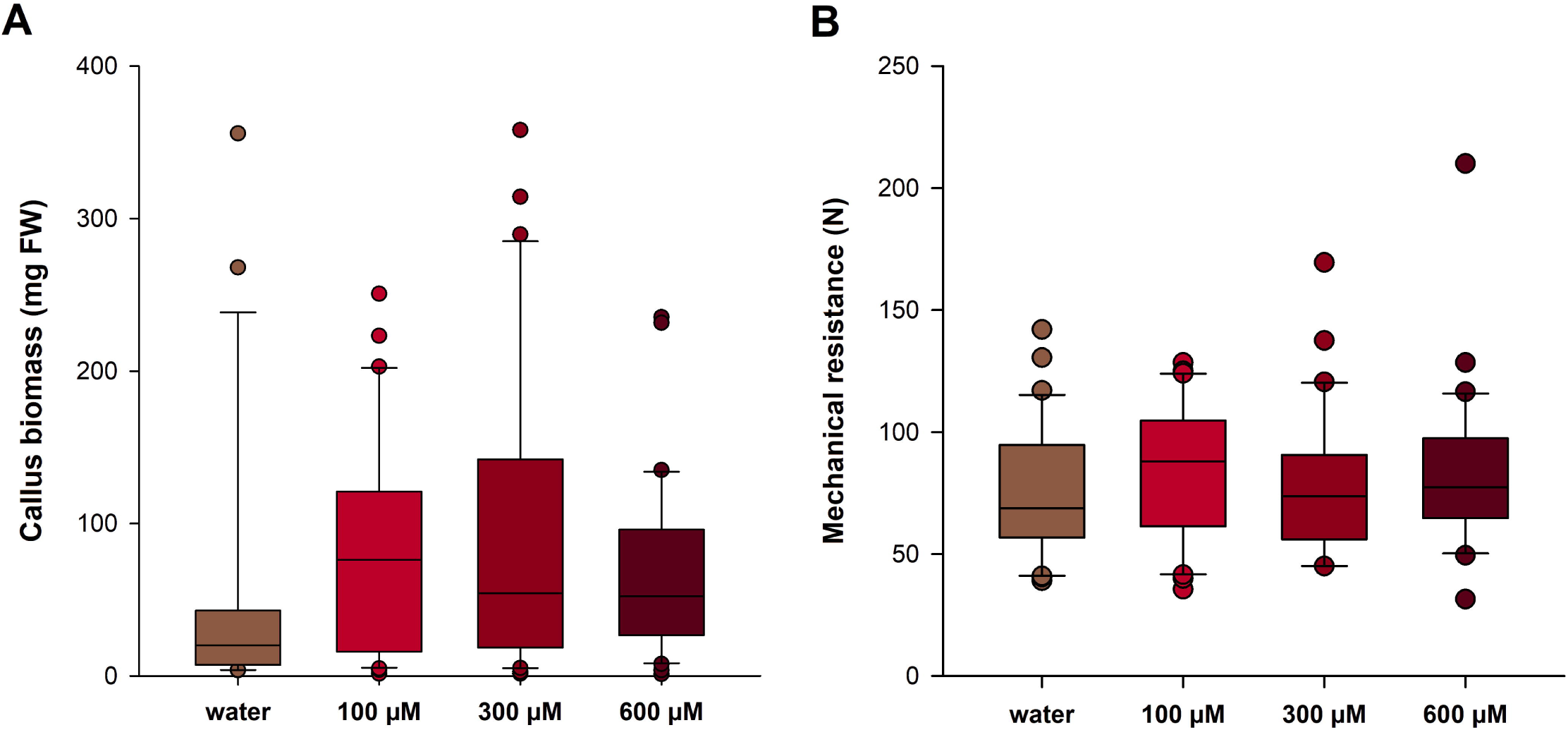
The effect of methyl jasmonate on A) callus biomass and B) the mechanical resistance of the graft interface 6 weeks after grafting. ALT TEXT: Box plots of A) callus biomass and B) the mechanical resistance of the graft interface of grafted treated with either water or three concentrations of methyl jasmonate.

## Discussion

### Identification of QTLs for component traits underlying graft union formation highlights the value of trait decomposition in genetic studies

Identifying QTLs of grafting-related traits has proved challenging for perennial fruit crops (Morel *et al*., 2026; Munoz *et al*., 2026; Pina *et al*., 2021). In this study, we were also unable to detect QTLs of grafting success rate despite studying up to 271 genotypes in one experiment. This could be due to different factors, but one limit in our and previous work has been the limited number of grafts performed for each genotype. As mentioned in the Introduction, it can take many years to have enough plant material to do a large number of grafts in perennial fruit crops, which can limit the possibility to do a large number of grafts per genotype. It is also a logistical challenge to graft so many scion/rootstock combinations with a high number of repetitions and it has been shown that non-genetic batch effects can dominate grafting success rates in large scale grafting experiments (Munoz *et al*., 2026). Furthermore, grafting success rate is multifactorial: it depends on the source of the plant material used for grafting, the skill of the operator, management practices and environmental conditions (from the initial callusing conditions to subsequent plant growth) (Hartmann, 2002). For these reasons, we decided to dissect grafting success rates into the grafting-related component traits of callus formation at the graft interface, mechanical resistance of the graft interface, shoot biomass and root number as well as callogenesis capacity on isolated stem internode segments. We successfully identified QTL for some of these grafting-related component traits; the heritability of these traits was low to moderate (from 0.29 to 0.5) suggesting that some of the variation in these traits is explained genetic factors. Whereas, the heritability of callus formation on isolated stem internode segments was moderate to high (0.51 and 0.71), suggesting that this trait is largely genetically determined. Similar levels of broad sense heritability have been found for callus induction on poplar leaves (Tuskan *et al*., 2018) and barley seeds (Benlioglu *et al*., 2025) suggesting that callus induction is genetically determined across a range of plant species and induction conditions. Despite the moderate to high levels of heritability for callus induction on isolated stem internode segments, we did not detect any significant QTLs for this trait. This could suggest that this trait is highly polygenic or that the population size was too small to detect any QTLs for this trait.

Broadly speaking, many of the grafting related traits and success rates were correlated with each other. This appears logical in that a plant with a well-developed graft union would be expected to have a certain amount of callus tissue and high mechanical resistance, which would trigger vigorous shoot and root growth. Interestingly, both the mechanical resistance of the graft union and shoot biomass were correlated with grafting success rates when measured 120 d (G) and 250 d (F) after grafting, suggesting that these could be a good proxies for phenotyping graft union development. It would be interesting to identify what determines mechanical strength of the graft union, it is generally assumed to be related to the formation of xylem vessels and vascular connectivity (Thomas *et al*., 2023), but this has just to be robustly tested. As attachment precedes vascular reconnection, it is possible that graft adhesion strength is also related to cell wall modifications in addition to vascular connections (Notaguchi *et al*., 2020).

Callus formation on isolated stem internode segments was correlated with callus formation at the graft interface only when considering dry mass, but with a low Pearson correlation coefficient. This suggests that the ability of a genotype to produce callus on isolated stem internode segments is somewhat unrelated to callogenesis at the graft interface. However, we did find that callus formation on isolated stem internode segments was correlated with callus fresh and dry mass suggesting that callus area measurements give a good indication of callogenesis capacity.

### Combining genetics, genomics and gene silencing to identify regulators of grafting

We identified 33 transcriptional regulators present within the confidence intervals of the QTLs of grafting-related component traits, which were significantly up-regulated in response to grafting in both the scion and rootstock. They included seven *ERF*, five *LOB*, three *bHLH*, four *GRAS* (including *PAT3*), *GATA8*, four *MYB* or *MYB-RELATED*, one *NAC* and two *WRKY* (including *WRKY65*). Orthologues of many of these transcriptional regulators are known to regulate wounding and stress responses, callus formation or grafting success in other plant species. For example, the orthologue of *PAT3/Vitvi10g00271* is *SCL13/AT4G17230*, which is part of the *PAT1* gene family, which has been shown to regulated callus formation and grafting success in both Arabidopsis and spruce (Feng *et al*., 2024b). We found that exogenous application of dsRNAs targeting *PAT3* reduced callus formation on isolated stem internode segments suggesting that this role is conserved in grapevine. In addition to *PAT3*, *PAT1/Vitvi19g00392* is in the confidence interval of the QTL on LG19 and is up-regulated in response to grafting but only in the scion. PAT1 in grapevine has been shown to interact with INDETERMINATE-DOMAIN 3/Vitvi12g00252 and regulate jasmonate levels via binding to LOX3/Vitvi09g00085 during cold treatment (Wang *et al*., 2021). *INDETERMINATE-DOMAIN 3* was not differentially expressed in response to wounding or grafting in grapevine, but *LOX3/Vitvi09g00085* was strongly up-regulated in response to grafting both scions and rootstocks (Loupit *et al*., 2025), and this gene is also located within the QTL on LG9. Hydroxy jasmonoyl isoleucine, an active jasmonate signal, has also been shown to accumulate at the graft interface of grapevine (Loupit *et al*., 2025). The role of jasmonate signals in regulated graft union formation is also supported by our observation that exogenous methyl jasmonate application increases callus development at the graft interface. As methyl jasmonate application did not affect the mechanical strength of the graft union, this potentially suggests that methyl jasmonate promotes callus proliferation, but not vascular differentiation.

Hormones have long been known to be important regulators of graft union formation in plants (Nanda & Melnyk, 2018). In grapevine, the exogenous application of auxins and cytokinins have been extensively studied in grapevine (Kose & Guleryuz, 2006) and a synthetic auxin in often used in commercial grafting waxes. There have been far fewer studies into the role of abscisic and jasmonic acids in graft union formation, however callus induction has been shown to be stimulated in Arabidopsis mutants of both jasmonic and abscisic acid synthesis and signaling (Ikeuchi *et al*., 2017). We also identified three signal peptides from the category non-cysteine-rich-peptides within our QTLs and up-regulated by grafting, wounding responses are well known to include signal peptide signaling such as systemin (Chen *et al*., 2020) and the recently identified REF1 (Yang *et al*., 2024), but the role of signal peptides in graft union formation remains to be studied.

Seven ERF TFs were up-regulated in response to grafting and within the QTLs of grafting-related traits, many ERF TFs have roles in wounding, grafting and regeneration competence, particularly those of the X subgroup (*ERF108-115*; Heyman *et al*. (2018)). Three ERF subgroup X TFs were up-regulated in response to grafting were found within the confidence interval of the QTL on LG7. The four other ERFs were one orthologue of *TARGET OF EARLY ACTIVATION TAGGED 2* (*TOE2/Vitvi13g00529*), which regulates plant antiviral immunity (Jiao *et al*., 2025), one and two from the subgroups I and VIII respectively. Because ERF114 and ERF115 have been shown to heterodimerize with GRAS TFs of the PAT1 family (Heyman *et al*., 2016), we attempted to silence both *PAT3* and an *ERF114* by the exogenous application of dsRNAs. However, we did not observe an additional inhibition of callus formation by the addition of dsRNAs against *ERF114*, suggesting that other subgroup X ERF TFs may interact with PAT3 in grapevine.

Four MYB TFs were up-regulated in response to grafting and located with QTL of grafting-related traits, one of them, *MYB102/Vitvi19g00306*, was identified as a potential hub gene regulating grafting responses (Loupit *et al*., 2025). In Arabidopsis, MYB102 is involved in hormone signaling and wounding responses (Denekamp & Smeekens, 2003). One MYB TF was a *MYB93* (*Vitvi07g02075*), in Arabidopsis MYB93 regulates lateral root formation and the expression of cell wall related genes (Uemura *et al*., 2023). And one of the other MYB TFs, *MYB148/Vitvi07g03055* is an orthologue of the *MYB68* subclade, which activate suberin biosynthesis and polymerization, while repressing cork cambium proliferation in Arabidopsis (Molina *et al*., 2025).

*Vitvi10g00618/WRKY65* was found to be up-regulated in both the scion and rootstock in response to grafting and was also within the QTL of callus biomass at the graft interface on LG10. In *V. amurensis*, WRKY65 is involved in ROS signaling and cold tolerance, and activates the expression of *PEROXIDASE 36/POX36/Vitvi04g01378* (Meng *et al*., 2025), but *POX36* expression was not up-regulated in response to grafting (Loupit *et al*., 2025).

Three *LBD11* genes were within the QTL of callus biomass at the graft interface on LG13 and highly up-regulated in response to wounding and grafting. The Arabidopsis orthologue is *AT2G28500* and this gene has been shown to regulate vascular cambium activity in interaction with ROS signaling (Dang *et al*., 2023). In Arabidopsis, LBD11 activates the expression of *POX71*, and *RESPIRATORY BURST OXIDASE HOMOLOGS D* and *F* (Dang *et al*., 2023), however, the grapevine orthologues of these genes were either not expressed at all or not up-regulated in response to grafting (Loupit *et al*., 2025).

Two of the *bHLH* TFs up-regulated in response to grafting and within the QTL of grafting-related traits on LG7 are orthologues of *NAI1*, which regulates the formation of endoplasmic reticulum bodies in Arabidopsis (Matsushima *et al*., 2003). *NAI1* genes have been previously identified as being particular highly expressed at the graft interface of heterografts (made of two different genotypes) compared to homo-grafts (a genotype grafted with itself) (Loupit *et al*., 2025).

One GATA TF was up-regulated in response to grafting and was within the confidence interval of the QTL for callus biomass at the graft interface on LG13. The pecan orthologue of *GATA8/Vitvi13g00614* has been identified as being highly expressed in response to grafting and silencing of this gene suggests that it has a role in defense responses (Zhang *et al*., 2025).

In summary, the 33 transcriptional regulators up-regulated in response to grafting and located within the confidence intervals of QTLs related to grafting included many orthologues of TFs known to regulate defense, wounding and regeneration in other plant species. The hypothesis that processes regulating graft union formation are conserved between distantly related species is support by the idea that interfamily grafts can be compatible at least in the short term, including grafts between grapevine and tobacco (Notaguchi *et al*., 2020). The pivotal role of jasmonate in regulating wounding responses was underlined by the five jasmonate related genes located within the QTLs and up-regulated in response to grafting and stimulation of callus formation at the graft interface by exogenous methyl jasmonate treatments. In addition, recently it has been shown that pre-treating melon scions and rootstocks with methyl jasmonate accelerates graft union formation (Zhu *et al*., 2025).

## Conclusion

Characterizing the genetic architecture of grafting success has proven to be challenging in many studies including this work. To overcome this challenge, we developed assays for grafting-related component traits such as quantifying the mechanical resistance of the graft union, callogenesis and post-grafting growth, and we were able to detect six QTLs significant at the whole genome level and 13 at the chromosome level. By cross-referencing the genes present within the confidence intervals with our previously published transcriptomic data, we identified 33 transcriptional regulators potentially underlying genetic variation in grafting-related traits. Orthologues of many of these regulators have roles in defense, wounding, jasmonate signaling, grafting and regeneration responses in other plant species. Silencing one of these TFs, *PAT3*, by the application of dsRNAs reduced callus development, while exogenous application of methyl jasmonate increased callus development suggesting that the regulation of grafting-related traits shares some regulatory features common across diverse plant species.

## Supplementary data

Table S1: Grafting success rates for all individuals studied and those individuals common to all years.

Table S2: Number of genotypes use in each experiment and common genotypes across different experiments.

Table S3. Characteristics of genes within QTL confidence intervals.

Table S4. Primer and dsRNA sequences

Table S5. Expression of genes within confidence intervals of QTLs and up-regulated in response to grafting.

Fig. S1: Heatmap of grafting success rates for 78 individuals in 2021, 2022 and 2024.

Fig. S2: Heatmaps of expression levels of cell wall and hormone related genes in response to grafting. Fig. S3: Correlation between callus surface area and callus fresh weight

## Acknowledgements

Thanks to UE 1442, UE Vigne, Unité Expérimentale Vigne & Vin Bordeaux, INRAE for the nursery. Thanks to the interns for their participation in the experiments, Hugo Doussy, Juliette Silvadier, Rose Touchard and Tarek El Mestari.

## Author contributions

SJC and EM: conceptualization, supervision and funding acquisition; MC, JPT, CH, ML and NH: methodology and investigation; MC and MM: formal analysis, validation and visualization, ML: data curation; SJC and ML: writing - original draft; all authors: writing - review and editing.

## Data availability

The data and photographs from this article are publically available in French government data repository, Recherche Data Gouv, https://doi.org/10.57745/GYCXEG.

## Abbreviations

BER: Vitis berlandieri 10585
CS: Cabernet-Sauvignon
dsRNA: double-stranded RNA
ERF/AP2: ethylene response factor/apetala2
F: field
G: greenhouse
GFP: green fluorescent protein
LG: linkage group
LOD: logarithm of the odds
LOX: lipoxygenase
PAT: phytochrome A signal transduction
POX: peroxidase
QTLs: quantitative trait loci
RUP: Vitis rupestris cv. Martin
TF: transcription factor
TMM: trimmed mean of M component
TOE: target of early activation tagged
WIND1: wound induced dedifferentiation1

